# A novel drug-inducible sex-separation technique for insects

**DOI:** 10.1101/2019.12.13.875716

**Authors:** Nikolay P. Kandul, Junru Liu, Alexander D. Hsu, Bruce A. Hay, Omar S. Akbari

## Abstract

Large sterile male releases are the gold standard for most insect population control methods and thus precise sex sorting is essential to the success of these technologies. Sex sorting is especially important for mosquito control because female mosquitoes bite and transmit diseases. However, current methods for insect sex sorting have deficiencies as they are error prone, low throughput, expensive, reduce male fitness, or lack cross species adaptability. Here we describe a novel drug-inducible system for insect sex-separation that demonstrates proof-of-principle for positive sex selection in *D. melanogaster*. The system exploits the toxicity of commonly used broad-spectrum antibiotics geneticin and puromycin and rescues only one sex. Sex specific rescue is achieved by inserting the sex-specific introns, *TraF* and *DsxM*, into the coding sequence of antibiotic resistance genes, *NeoR* or *PuroR*. We engineer a dual sex-sorter gene cassette and demonstrate sex specific, constitutive expression of NeoR and PuroR proteins in females and males, respectively. When raised on geneticin supplements, this sex-sorter line established 100% positive selection for female progeny, while the food supplemented with puromycin generated 100% male progeny. This system is 100% efficient and operates at remarkably low fitness costs in *D. melanogaster*. Since the described system exploits a conserved sex-specific splicing mechanism and reagents, which are active in many insects, it has the potential to be adaptable to insect species of medical and agricultural importance.

**GRAPHIC ABSTRACT:** 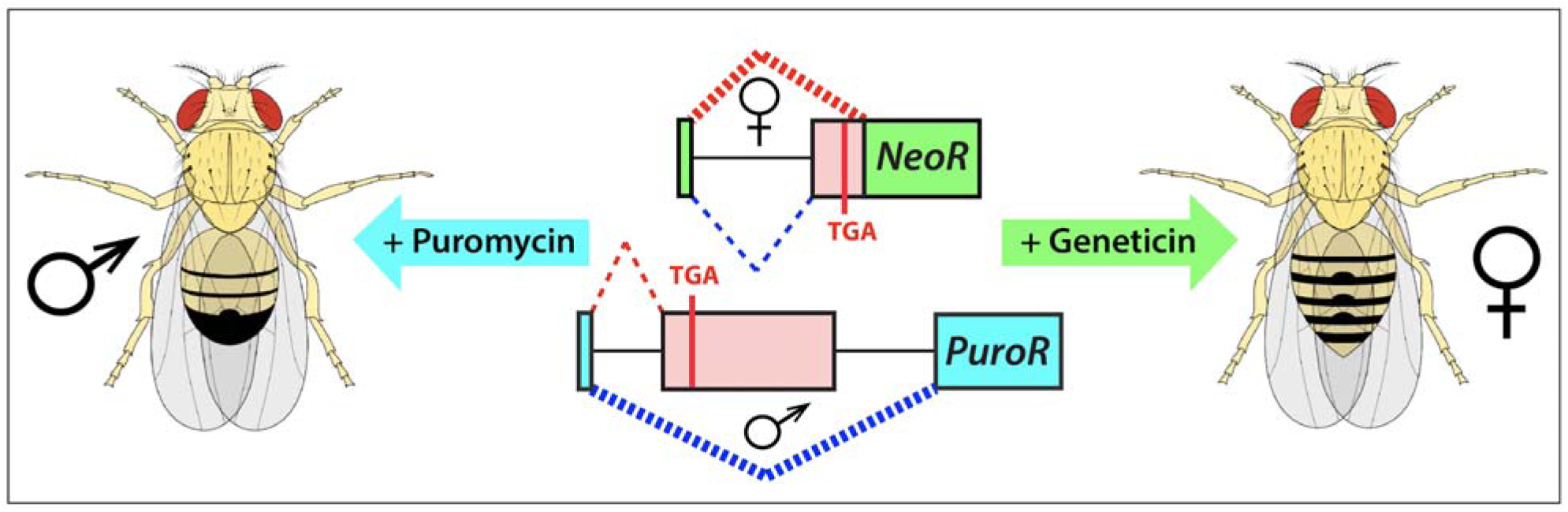

## INTRODUCTION

The ability to generate and release large numbers of males that are not able to sire progeny into natural populations is a key requirement for many current approaches in insect population control. In 1955, Edward Knipling proposed releasing sterile males to suppress insect populations in so called Sterile Insect Technique (SIT)^1^. Since then, SIT was successfully implemented to eradicate *Cochliomyia hominivorax*, the new world screw-worm fly, in the U.S. and Mexico^2^ and suppress populations in a variety of other insects^3,4^. However, Knipling’s goal of sexing sterilized insects by removing females prior to their release was challenging to accomplish even in *C. hominivorax* and has limited the implementation of SIT to agricultural pests. Field trials and models clearly showed that releasing only sterile males, which were not distracted by accompanying females, significantly improves the efficiency of population suppression and can result in significant cost reduction^1,5^. Other related methods of insect control, such as Release of Insects carrying a Dominant Lethal (RIDL)^6^ and the *Wolbachia*-mediated Incompatible Insect Technique (IIT)^7–9^, also require consistent sexing methods to avoid female releases. Notably, IIT programs are based on the release of *Wolbachia* infected males incompatible with the wild type females, which are free of a *Wolbachia* strain, and the accidental release of only a small proportion of *Wolbachia* infected fertile females can immunize a target population against the *Wolbachia* strain used in a particular IIT program. Furthermore, since mosquito females are blood feeders and transmit many diseases, a reliable sexing method to guarantee female elimination among released insects is highly desirable for the implementation of SIT, RIDL, or IIT programs in mosquito species.

Over the years, a diverse array of sexing methods has been developed for several insect species. First, mechanical separation makes use of physical differences between the two sexes, which may include morphology (size^10^ and shape), coloration, hatch timing, and behavioral differences (female blood feeding^11^ and male swarming)^12–15^. Second, various genetic approaches have been developed. In a classic Genetic Sex Separation (GSS) approach, a conditional lethal, a resistance to dieldrin insecticide (*Rdl*), was translocated on the Y chromosome through irradiation-induced chromosomal rearrangements in multiple *Anopheles* species allowing for the permissive survival of only males when exposed to dieldrin^16–19^. Another GSS approach relies on generating transgenic insects carrying fluorescent markers either genetically linked to sex chromosomes^19–22^ or with sex-specific expression^23,24^ that can be mechanically sorted. Finally, sex separation can also be achieved by negative selection against females using conditional sex-specific lethal transgenes that are repressed by continuous feeding of tetracycline (a Tet-Off system). The sex-specificity of markers’ expression and conditional lethal transgenes was generated serendipitously^6,25,26^ or by incorporation of a female-specific intron into a coding sequence of a lethal transgene^27–30^.

Notwithstanding, the current methods for sex-sorting of insects have many shortcomings and are not easily transferable to other species. The methods based on mechanical separation of different sexes are entirely species specific and cannot be adopted to other species. In addition, they are not practical in most insects, since they require optimal rearing conditions, lead to a significant error rate, and are not suitable for high-throughput insect production^13,15,25,31,32^. Classic GSS approaches were achieved serendipitously in one species at a time, and thus are species specific. Fluorescent sex sorting requires that each larva is examined and sorted individually, and as a result this approach is generally not suitable for large-scale programs demanding a high throughput production. Furthermore, induced chromosomal rearrangements and sex chromosomal linkage are genetically unstable, and would frequently break down as a result of chromosomal recombination when large numbers of insects were raised^15,25,31,33^. In the Tet-Off approaches, tetracycline must be continually supplied to prevent lethality during mass rearing. Consequently, this constant exposure can generate unwanted fitness effects such as negative effects on mitochondrial functions^34–36^. Therefore, to improve the efficiency of current methods for insect population control, novel approaches for sex-sorting that are genetically stable, suitable for a large-scale insect production, and can be adopted to different insect species are required^13,15,32^.

Here we describe a positive drug-inducible GSS system for insects and demonstrate its proof-of-principle in *Drosophila melanogaster*. Two genes confirming resistance to specific drugs are expressed in opposite sexes by incorporating sex-specific introns into the coding sequence of drug-resistance genes. In the absence of sex-selection, the transgenic strain harboring a sex-sorting gene cassette is maintained on normal food. When the insects of a particular gender are required, the transgenic strain is raised on the food supplemented with the corresponding drug. Members of the sex selected against are not resistant to a selecting drug and are killed early during development, as the first instar larvae, resulting in emergence of 100% adults of the desired gender. The described GSS system is resistant to genetic recombination, chromosomal rearrangements and other mutations; and it can be potentially adapted to different insect species with known sex-specific introns.

## RESULTS

To engineer a drug-inducible sex-selection system in *D. melanogaster* we utilized two common antibiotic-resistance genes, *puromycin N-acetyltransferase* (*PuroR)* and *aminoglycoside phosphotransferase* (*NeoR)*, previously demonstrated to confer resistance in eukaryotic cells, *D. melanogaster* S2 cells, and *D. melanogaster* larvae, to corresponding water soluble antibiotics, puromycin and geneticin, respectively^37–40^. To determine the toxic doses for these drugs in *D. melanogaster*, we raised wild type fly larvae on food supplemented with either increasing concentrations of puromycin or geneticin (0 mg/ml, 0.2 mg/ml, 0.4 mg/ml). From this experiment, we determined that a concentration of 0.2 mg/ml of either drug was quite toxic, however it permitted survival of some *D. melanogaster* larvae to adulthood (2.2% ± 1.5% for puromycin; and 14.0% ± 6.5% for geneticin, Table S1), while concentrations above 0.4 mg/ml completely inhibited development with nearly all larvae unable to mature past the 1^st^ instar stage, and 100% of larvae perishing before adulthood on the supplemented food (for each treatment, expected n > 500 found 0; replicates (N) = 5; *P* < 0.001: Figure S1a, Table S1).

Following confirmation of the toxic doses, we then tested if we could rescue this toxicity by transgenic expression of antibiotic resistance genes *PuroR* or *NeoR* integrated into the *D. melanogaster* genome. We engineered a PiggyBac (PB) transposable element that encoded a constitutive baculovirus promoter *Hr5IE1* driving expression of dsRed as a selectable marker (*Hr5IE1-dsRED*)^41^. To provide continuous and ample supply of an antibiotic resistance protein, we utilized a strong ubiquitous baculovirus promoter *Opie2*^*42*^ to express the *PuroR* or *NeoR* gene (*Opie2-PuroR* or *Opia2-NeoR*) and inserted the gene cassette in an opposite orientation relative to *Hr5IE1-dsRED* to avoid any transcriptional read through effects (Figure 1a). We generated a few transgenic lines harboring one copy of either *Opie2-PuroR* or *Opie2-NeoR* and let them lay eggs on fly food supplemented with either puromycin or geneticin, respectively. Interestingly, when non-balanced transgenic fly lines, which contained both transgenic and wild type (*wt*) flies, were raised on food supplemented with either puromycin (0.4 mg/ml) or geneticin (0.4 mg/mL), only transgenic flies carrying a single copy of the resistance gene matched to a supplemented drug emerged, while both *wt* and non-matched transgenic larvae perished (for each treatment, n > 500; N > 6; *P* < 0.001: Figures 1a–b, S1b; Table S1). Taken together these results strongly indicated that baculovirus promoter driven expression of *PuroR* and *NeoR* genes was able to dominantly rescue the larval lethality caused by consumption of toxic doses of either puromycin or geneticin (Figure 1a–b).

**Figure 1.**
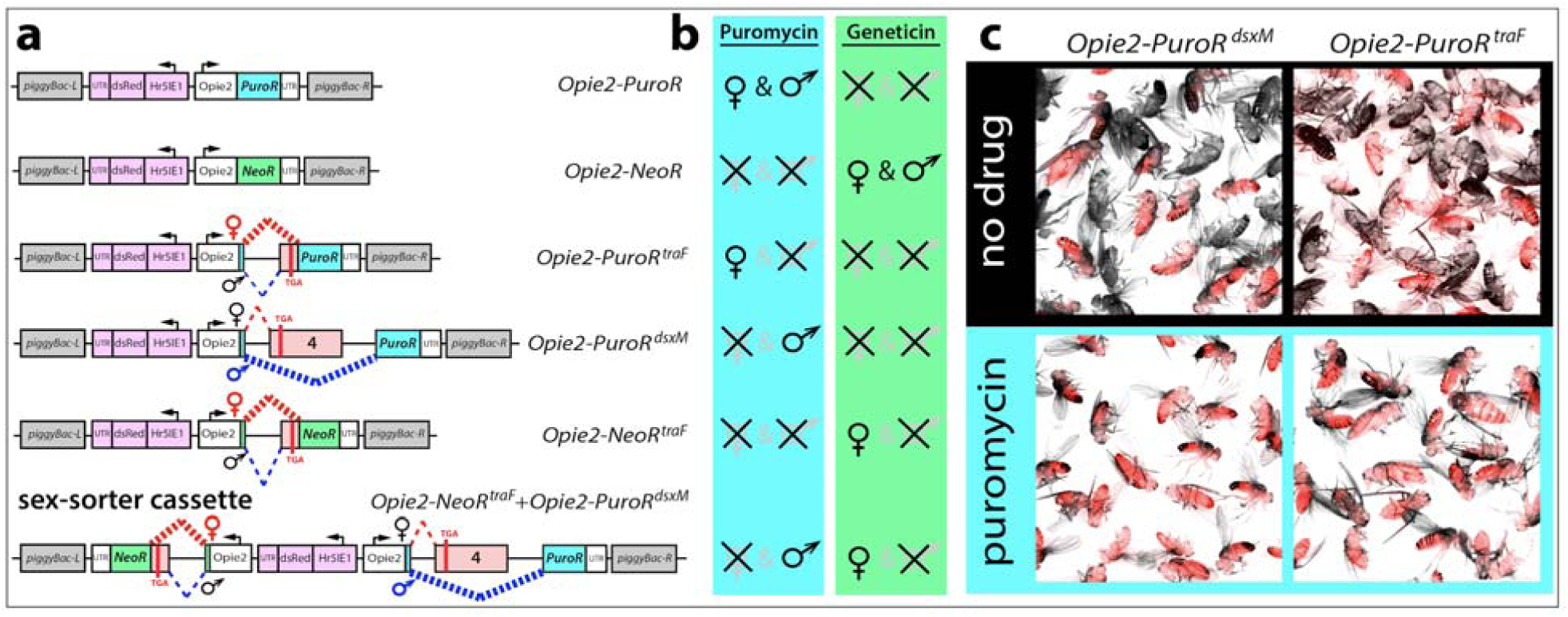
Development of sex-sorter cassette in *Drosophila*. (a) Schematic maps of genetic constructs engineered and tested in the study. (b) Expression of antibiotic resistance genes (*PuroR* and *NeoR*) throughout *Drosophila* development confers resistance to puromycin and geneticin, respectively, supplemented on fly food. To insure that functional antibiotic-resistance proteins will be produced only in one or the other gender, sex-specific introns from two sex-determination genes (*tra* and *dsx*) were inserted into coding sequences of *PuroR* and *NeoR*. The entire sequences of female-specific *traF* and male-specific *dsxM* introns (highlighted in pink) are spliced out in the corresponding sex, but some sequences carrying a stop codon (TGA) are retained in the opposite sex (Figure S2). The transgenic flies harboring one copy of a genetic construct were identified by a strong ubiquitous expression of *dsRed* (highlighted in purple). (c) Expression of *Opie2-PuroR*^*dsxM*^ or *Opie2-PuroR*^*traF*^ transgene rescues only transgenic males or females (red fluorescence) raised on the food supplemented with puromycin, while all wild type flies (no red fluorescence) and the transgenic flies of selected-out sex die during early development.

We next determined whether we could exploit the endogenous sex determination machinery to promote antibiotic selection in a sex specific manner. In *D. melanogaster*, the female-specific intron between *transformer* (*tra*) exons 1 and 2 (*traF*) is spliced out entirely in females, while in males some sequence remains producing a stop codon resulting in premature termination of Tra protein^43^ (Figure S2). Inversely, the male-specific intron between *double sex* (*dsx*) exons 3 and 5 (*dsxM*) is spliced out in males, while in females only its small part is spliced out and an entire exon 4 carrying a premature stop codon remains^44^ (Figure S2). Given these known intronic splicing patterns, we hypothesized that if we insert these sex-specific introns into the coding sequences of *PuroR* or *NeoR*, functional proteins would only be produced in one or the other gender leading to its survival, while the opposite gender would perish.

To test this hypothesis, we generated plasmids encoding previously characterized sex-specific introns *traF*^*43*^ or *dsxM*^*44*^ within the coding sequence of *PuroR*, generating two types of synthetic transgenes, *Opie2-PuroR*^*traF*^ and *Opie2-PuroR*^*dsxM*^ (Figure 1a). We then made transgenic flies harboring one copy of either *Opie2-PuroR*^*traF*^ or *Opie2-PuroR*^*dsxM*^ and let them develop on fly food supplemented with puromycin. Because transgene integration site can affect gene expression and sex-specific splicing^45^, a few heterozygous transgenic lines were assessed for each construct. Raising two independent lines carrying one copy of *Opie2-PuroR*^*traF*^ on food supplemented with puromycin at 0.4 mg/mL indeed resulted in emergence of only female flies (n = 421, N = 10, *P* < 0.001: Figures 1a–b, S1c); while three *Opie2-PuroR*^*dsxM*^ lines raised on the same fly food gave rise to male-biased progeny (n = 741, N = 13, *P* < 0.001: Figure S1c); and one of them produced only male progeny on the higher puromycin concentration, 1.0 ml/mL (n = 81, N = 3, *P* < 0.001: Figures 1a–b, S1c; Table S1). Notably, when we grew the mixture of highly selective *Opie2-PuroR*^*traF*^ or *Opie2-PuroR*^*dsxM*^ transgenic line and *wt* flies on food supplemented with 1.0 of puromycin, only female or male transgenic flies marked with dsRed fluorescence emerged while no *wt* larvae survived to adulthood (Figure 1c). To determine the versatility of the system and to establish a geneticin-mediated sex-selection system, we inserted the *traF* intron into the coding sequence of *NeoR* and generated the transgenic flies carrying *Opie2-NeoR*^*traF*^. Three independent heterozygous *Opie2-NeoR*^*traF*^ lines raised on food supplemented with geneticin at 0.4 mg/mL resulted in female-biased progeny (n = 620, N = 9, *P* ≤ 0.05: Figure S1c); and one of them enforced 100% female selection on the higher geneticin concentration, 1.0 ml/mL (n = 188, N = 3, *P* < 0.001: Figures 1a–b, S1c; Table S1). Taken together these results indicate that by inserting the *traF* or *dsxM* sex-specific introns into the coding sequence of *PuroR* or *NeoR* the expression of functional antibiotic resistance genes can be limited to one or the other sex of *D. melanogaster*.

To enable positive drug selection of either sex from a single synthetic construct (termed a sex-sorter cassette from hereon), we tested if we could combine the two separate sex-selection systems together (*Opie2-NeoR*^*traF*^ + *Opie2-PuroR*^*dsxM*^: Figure 1a). We engineered a sex-sorter cassette plasmid, generated three independent transgenic lines harboring the cassette, and tested them by raising heterozygous flies on food supplemented with either puromycin or geneticin at 0.4 and 1.0 mg/mL. For two of three tested lines, only female flies emerged on food supplemented with 1.0 mg/mL of geneticin (n = 335, N = 6, *P* < 0.001); and only males were recovered from vials containing 1.0 mg/mL of puromycin (n = 210, N = 6, *P* < 0.001: Table S1), while the lower drug concentration was not sufficient to enforce emergence of single-sex progeny for each of three tested transgenic lines harboring a single copy of the sex-sorter cassette (Figure S1c). These results strongly indicate that by raising flies harboring one copy of the sex-sorter cassette on the food supplemented with either drug at 1.0 mg/mL we can dominantly control which gender survives to adulthood.

We next explored an opportunity to lower the drug concentration and still enforce 100% efficient sex sorting by doubling a copy number of the sex-sorting cassette. The homozygous sex-sorter line established from the flies carrying *Opie2-NeoR*^*traF*^+*Opie2-PuroR*^*dsxM*^ cassette integrated on III chromosome (line #3 on Figure S1c) did not manifest any obvious fitness defect, and was used for further analysis. We raised the homozygous flies on food supplemented with either puromycin or geneticin, titrating concentrations down from 1.2 mg/mL, and quantified gender percentages of emerging flies. We discovered that the addition of puromycin at the final concentrations of 1.2, 1.0, 0.8, 0.6, and 0.4 mg/mL resulted in 100% male progeny. Even 0.2 mg/mL of puromycin caused a significant increase in the male ratio, from 43.0% ± 1.3% vs 62.7% ± 2.15% (N = 3, *P* ≤ 0.001; Figure 2a). Inversely, supplementing food with geneticin to the final concentrations of 1.2, 1.0, 0.8, 0.6, 0.4, and 0.2 led to 100% female progeny. For geneticin, even 0.1 mg/mL led to a significant increase of female progeny, from 56.1% ± 1.1% to 78.0% ± 4.6% (N = 3, *P* ≤ 0.01; Figure 2b; Table S2). Importantly, the antibiotic-mediated selection of fly gender does not affect fertility of recovered flies. We repeatedly tested the fertility of males and females with two copies of the sex-sorter cassette raised on the highest tested concentration of puromycin or geneticin (1.2 mg/mL), respectively, by housing them with *wt* virgin flies of an opposite sex on non-supplemented food. In each case, numerous progeny containing both genders without any obvious phenotypic defects were obtained. Taken together, these results show that flies of a specific gender, can be generated with 100% efficiency by simply raising the flies on food supplemented with either puromycin (male selection) or geneticin (female selection) at 0.4 mg/mL.

**Figure 2.**
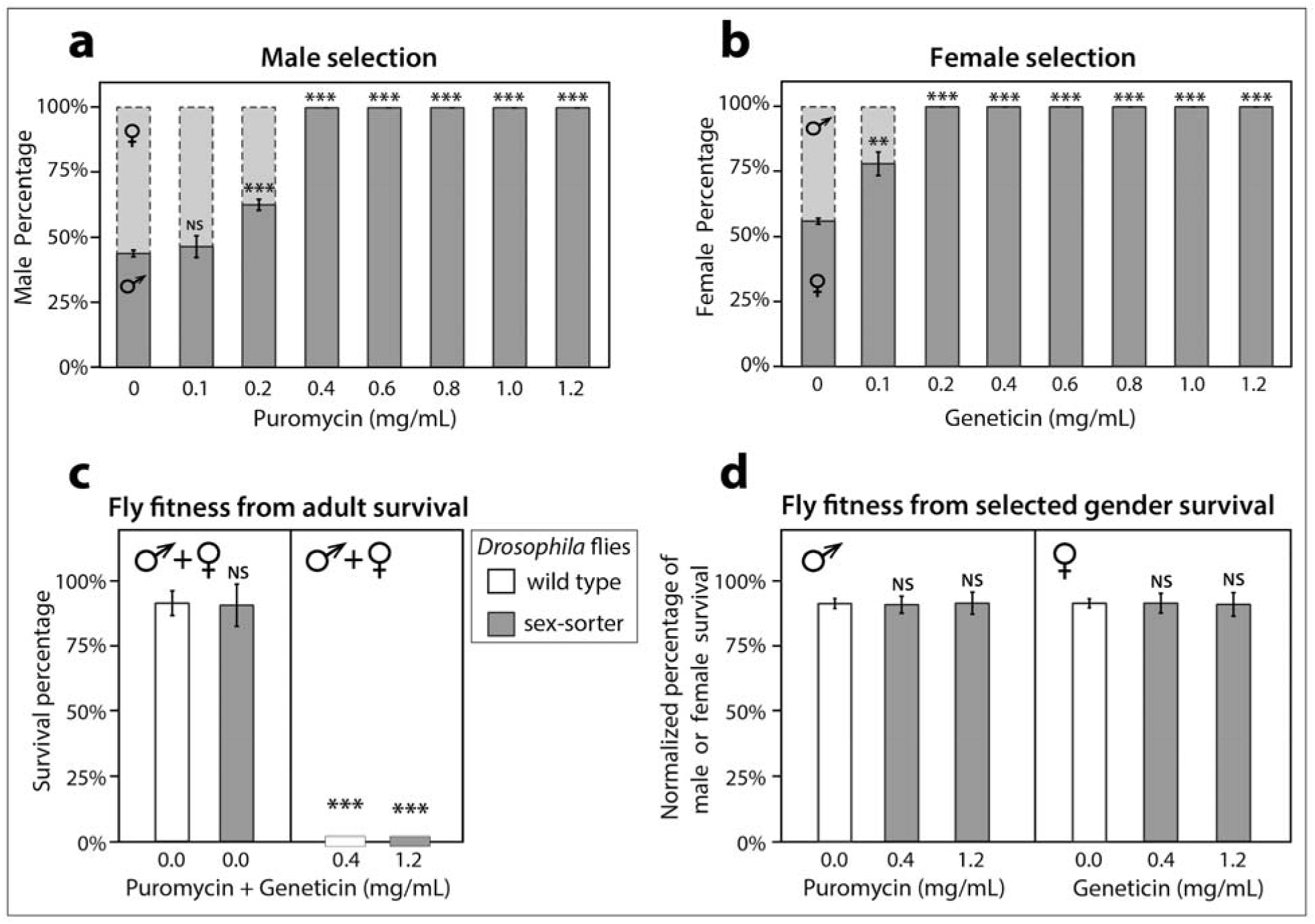
Sex selection and fitness of the flies carrying two copy of sex-sorter cassette. The sex-sorter cassette includes *NeoR*^*traF*^ and *PuroR*^*dsxM*^ genes that confer resistance to puromycin and geneticin antibiotics, respectively, in the sex-specific manner (Figure 1a). The *PuroR*^*dsxM*^ gene is properly spliced and results in expression of the functional PuroR protein only in males, while *NeoR*^*traF*^ expresses the NeoR protein only in females. To estimate the lowest concentration of an antibiotic at which male or female selection is enforced at 100%, the homozygous sex-sorter flies were raised on various concentrations of puromycin or geneticin. (a) Only male flies emerged from the food supplemented with 0.4 mg/mL or more of puromycin. (b) Raising the same flies on the food containing 0.2 mg/mL or more of geneticin resulted in the emergence of only female flies. To compare the fitness of homozygous sex-sorter flies to that of wild type (*wt*) flies, the embryo-to-adult survival of both fly types were compared under normal and selective conditions. (c) Embryos of both sex-sorter (gray bars) and *wt* flies (white bars) survived to the adulthood equally well on food without any antibiotics and died on the food supplemented with both puromycin and geneticin to 0.4 and 1.2 mg/mL. (d) The survival of male or female sex-sorter flies under selection treatments was statistically identical to that of the corresponding gender from *wt* flies raised under normal conditions. Bar plots show the average ± SD over at least three biological replicates. Statistical significance was estimated using a *t* test with equal variance. (*P* ≥ 0.05^ns^, *P* < 0.05*, *P* < 0.01**, and *P* < 0.001***).

Two possible reasons can explain why the sex selection is not effective at lower drug concentrations. First, the concentration may be too low to enforce effective selection as *wt* flies can also survive at these lower concentrations. We previously found that *wt* flies cannot be raised on 0.4 mg/mL of either puromycin or geneticin: fly development arrests at the larval 1^st^ instar stage. To test whether *wt* flies can survive on concentrations lower than <0.4 mg/mL, we raised *wt* flies on 0.200, 0.100, 0.050, and 0.025 mg/mL of puromycin or geneticin. Notably, while development of flies raised on food supplemented with each antibiotic was delayed, a few flies repeatedly emerged on concentrations ≤ 0.2 mg/mL for each drug indicating that these concentrations are indeed too low for complete selection. Second, some antibiotics may degrade over time becoming insufficient to continue selecting against the opposite gender. In our experiments, we found that puromycin, was indeed unstable over time as we observed that after collecting exclusively male flies from the vials with 0.4 mg/mL of puromycin for eight straight days at +25°C, a few female flies would emerge starting at day 9 (Table S3). However, for geneticin at the same concentration (0.4 mg/mL), only female flies emerged from the vials supplemented with the antibiotic. This indicates that both *wt* fly survival and antibiotic degradation contribute to the lack of effective sex selection at the concentrations lower than < 0.4 mg/mL.

To estimate the fitness costs of two copies of the sex-sorter cassette to its carriers, we compared the fitness of homozygous sex-sorter flies to that of *wt* flies. We found that the survival rates, calculated as a percentage of embryos surviving to adults, for both lines raised on non-supplemented food, were not statistically different from *wt* flies: 91.2% ± 2.8% of *wt* embryos survived to adulthood vs 90.4% ± 8.1% of transgenic sex-sorter embryos survived to adulthood (N = 5, *P* = 0.85: Figure 2c). Moreover, supplementing fly food with both puromycin and geneticin at either 0.4 or 1.2 mg/mL totally arrests the life cycles of both fly lines, as expected since neither *wt* nor flies harboring two sex-sorter cassettes are resistant to both drugs simultaneously (Figure 2c). To further measure fitness, we compared the survival rates of each gender, normalized as a percentage of male or female embryos surviving to adults, compared between *wt* flies raised on food without any antibiotic and the homozygous sex-sorter flies raised on food supplemented with either puromycin or geneticin. The percentage of *wt* males that emerged on non-supplemented food was similar to that of sex-sorter males that emerged from the food supplemented with puromycin at concentrations of either 0.4 or 1.2 mg/mL: 91.2% ± 1.9% vs 90.6% ± 3.2% (N = 5, *P* = 0.93) and 91.2% ± 4.3% (N = 5, *P* = 0.77, Figure 2d). Similarly, the percentage of hatched *wt* females was not significantly different from sex-sorter females emerged on the food supplemented with geneticin at concentrations of either 0.4 or 1.2 mg/mL: 91.4% ± 1.7% vs 91.2% ± 2.8% (N = 5, *P* = 0.93) and 90.7% ± 4.5% (N = 5, *P* = 0.77, Figure 2d; Table S3). Finally, we also estimated the competitiveness of drug-sexed males with two copies of the sex-sorter cassette. We found that homozygous males raised on the food supplemented with 0.4 mg/mL of puromycin were as competitive as *wt* males at female mating and were able to sire half of female progeny in the presence of *wt* males (47.8 ± 8.6%, N = 10, progeny n = 1715, Table S4). These results indicate that the percentage of transgenic eggs successfully developing to viable male or female flies on drug-supplemented food is statistically as high as that of *wt* flies raised on non-supplemented food, indicating that neither the sex-sorter cassette nor the exposure to puromycin during larval development incurred significant costs on development and male courtship.

## DISCUSSION

We describe the proof-of-concept of a novel GSS approach in *Drosophila*. Its design is based on the sex-specific expression of two antibiotic-resistance genes (*NeoR*^*38*^ or *PuroR*^*37*^), which is accomplished by incorporation of introns that splice in sex-specific manner, *traF* and *dsxM*. Females or males are positively selected by rescuing their development on the food supplemented with either geneticin or puromycin. The described drug-inducible GSS approach has several advantages other traditional genetic and mechanical methods for insect sex-sorting: genetic stability, positive selection, portability across different insect species, low expenses for maintenance, low fitness cost, and adaptability for high throughput sex-sorting.

This system, unlike many GSS methods, is inherently resistant to breakage resulting from chromosomal translocations, recombination, and mutations. Any of these events causing a loss-of-function (LOF) mutation in *NeoR*^*38*^ or *PuroR*^*37*^ antibiotic resistance gene will be selected out since the flies harboring LOF mutations will not reproduce on the food supplemented with the corresponding antibiotic. Traditionally the construction of genetic sexing lines was based on linking a selectable marker gene, such as insecticide resistance^12,18,19^, the temperature-sensitive lethal (TSL), and phenotypic loci^25^ to a Y-autosome or X-autosome via induced chromosomal translocations. Such engineered lines are not stable, and chromosomal rearrangements will break the genetic linkage between a selective marker and a sex chromosome^13,15,18,25,31,32^. It was found that when large numbers of insects were produced from an engineered GSS line, rare chromosomal rearrangements persisted or were selected for, and would contaminate the original line making it unusable for sex sorting^15,32^. The novel approach does not require the genetic linkage to sex chromosomes, instead it relies on the sex-specific rescue of lethality and any mutations affecting the functionality of antibiotic resistance genes will be selected out during sex-sorting. To establish sex-specific splicing of the *NeoR* or *PuroR* genes in *Drosophila*, we inserted two endogenous introns from ultra-conserved genes in the sex determination pathway, *transformer* (*tra*) and *double sex* (*dsx*) (Figures 1a, S2) into coding sequences of *NeoR* and *PuroR*. The *traF* and *dsxM* intronic sequences are spliced out entirely only in females or males, respectively, and ensure that functional NeoR and PuroR proteins are expressed in one or the other gender. The positive selection of females or males by rescuing their lethality during development on the food supplemented with geneticin or puromycin guarantees that they harbor functional the sex-sorter gene cassette. The described method can be easily adapted to generate the large numbers of male or female insects.

The sex-sorter gene cassette was designed to be widely transferable between insect species. It was engineered in the *PiggyBac* transposable element, which has been shown to be portable across many insect species^46^. Puromycin and geneticin are general antibiotics that arrest growth in both prokaryotic and eukaryotic cells. The resistance genes for these antibiotics are well established and routinely used as selective markers for transgenesis of cell cultures, and therefore are expected to be effective and portable across a wide variety of insects^37–40^. Likewise, the *Opie2* and *Hr5IE1* regulatory sequences that promote expression of the sex-specific antibiotic resistance genes and the transformation marker, respectively, originate from a baculovirus known to infect a large variety of insects^41,47^. Most importantly, the underlying mechanism of how we achieve the sex-specific antibiotic expression via sex-specific alternative splicing is conserved across diverse insect species. For example, both the male-specific *dsx* and female-specific *tra* introns are conserved across many insects^48^, indicating that it should be straightforward to transfer this positive sex selection system into other insect species. In fact, the conserved female-specific *tra* intron has already been used to confer the female-specific lethality by splicing correctly a suicide gene, which is repressed by continuous tetracycline feeding or activated by a heat shock, in a few pest insect species^27–30^. While this system was demonstrated using the *PuroR* and *NeoR* antibiotic resistance genes, the concept of using sex specific introns as a positive sex selection approach may work for a wide range of other types of drug or insecticide resistance genes. These could include, for example, the cytochrome p450 genes that have been shown to confer resistance to a wide range of insecticides including DDT, nitenpyram and dicyclanil^49^.

In contrast to current genetic methods for sex sorting based on the repressible lethality (negative selection), the described design does not require continuous drug feeding for maintenance of the transgenic line^6^. Antibiotics, puromycin or geneticin, are supplied only to enforce the sex selection by rescuing the selected gender (positive selection) and ‘killing’ the opposite gender. Unlike Tet-Off systems with conditional lethal transgenes^6,27–29^, no antibiotic is required for survival and maintenance of the sex-sorter transgenic line. We demonstrate that drug selection occurs early on, at the first instar stage, and thereafter the surviving sex does not compete for resources with the opposite sex and can be maintained on a regular food. This transient exposure of antibiotics could further reduce any potential fitness costs and could also reduce costs associated with drug selection as smaller quantities could be used at only the early instar stages, a factor that may play a significant role in large-scale projects.

The sex-sorting gene cassette does not directly affect the fitness of its carriers. Unlike the GSS methods utilizing the negative selection^6,27–29^, the sex-sorter cassette does not include any toxic or suicide gene that could leak and affect the organismal fitness. We did not expect that the expression of PuroR and NeoR would be costly to the fly fitness, especially when only one antibiotic resistance protein expressed in each gender. Instead, the location of transgene integrations in the *Drosophila* genome could have a strong effect on the fitness of transgenic flies^45^, and we assessed multiple integration sites for each genetic construct (Figure S1). The homozygous sex-sorter transgenic line generated in the study harbors two copies of the sex-sorter cassette and have the same egg-to-adult survival rate for each gender as that in *wt* flies even when the transgenic flies were raised on food supplemented with antibiotics to enforce sex-selection (Figure 2d). We also found that the homozygous sex-sorter males raised on food supplemented with puromycin were able to compete effectively with the *wt* males for female mates and sired on average a half of the progeny. The competitiveness of sex-sorted males against *wt* males is of great importance^1,5^, since many insect control methods such as SIT, RIDL, and IIT rely on male releases. We think that the synthetic genetic circuit that relies on the positive, instead of negative, selection can ameliorate some of its negative effects of the organismal fitness.

The positive drug-inducible GSS system presented here complies with the seven key requirements for the efficient sex separation technology, referred as the “7 Ses”^13^. (1) Small, the gender selected against die early in development and does not compete for resources with the selected gender. (2) Simple, a required gender is positively selected by simply raising the transgenic insects on food supplemented with one or the other drug. (3) Switchable, the sex-sorter transgenic flies are maintained on a regular food. (4) Stable, the constitutive expression of functional antibiotic-resistance protein in sex-specific manner guarantees survival of a specific gender on drug-supplemented food, and any LOF mutations in the antibiotic-resistance gene are selected out. (5) Stringent, our data show that the sex sorting is enforced at 100%. (6) Sexy, the sex sorting happens genetically during insect development and does not required insect handling. It is mediated by the positive selection and its genetic circuit does not include any toxic or suicide genes resulting in the minimal fitness cost. (7) Sellable, since we describe the proof-of-concept in the *Drosophila* model species, we cannot assess whether this technology will be acceptable for a specific insect in a target release area. Nevertheless, the described GSS system can be potentially adapted to other insect species, especially in Dipteran order.

## MATERIALS AND METHODS

### Antibiotics and antibiotic resistance genes

Puromycin is a water soluble aminonucleoside antibiotic produced by the bacterium *Streptomyces alboniger*; it inhibits protein synthesis by disrupting peptide transfer on ribosomes causing premature chain termination during translation, and is thus a potent translational inhibitor in both prokaryotic and eukaryotic cells^50^. Geneticin (G418) is a water soluble aminoglycoside antibiotic produced by the bacterium *Micromonospora rhodorangea*; it interferes with 80S ribosomes and protein synthesis, and is therefore commonly used as a selective agent for eukaryotic cells (www.thermofisher.com). *PuroR* (*pac*)^37^ from *Streptomyces alboniger* encodes *puromycin N-acetyltransferase* and its expression in bacteria and mammalian cells confers resistance to puromycin. *NeoR* (*neo*)^38^ encodes *aminoglycoside phosphotransferase* and its expression in bacteria and mammalian cells confers resistance to geneticin, kanamycin, and Geneticin® (G418).

### Molecular Biology

To support transgenesis in diverse insect species, genetic constructs were built inside a PiggyBac (PB) JQ352761 plasmid^51^ digested with FseI and AvrII. The PB (formerly IFP2) transposon was originally defined in the *Trichoplusia ni* cabbage looper moth^52^ but has become a transposon of choice for genetic engineering of a wide variety of species, particularly insects^46^. We used Gibson assembly method to engineer genetic constructs. Protein sequences of *PuroR (pac)*^*53*^ and *NeoR* (*neo*)^38^ were back translated, codon optimized for *Drosophila* in Gene Designer 2.0 (https://www.dna20.com/resources/genedesigner), and synthesized as gene-blocks by Integrated DNA Technologies®. Both genes were expressed constitutively under *Opie2* regulatory sequences that originate from the baculovirus *Orgyia pseudotsugata* multicapsid nuclear polyhedrosis virus (OpMNPV)^42^ and amplified from JQ352760 plasmid^51^. The transformation marker dsRed, a red fluorescent protein, amplified from KC733875 plasmid^54^ was ubiquitously expressed under the *Hr5IE1* regulatory sequence amplified from KC991096 plasmid^55^. SV40 3’UTR from pAc5.1-V5-HisB (Invitrogen®) terminated transcription of transgenes. To confer sex-specific translation of *PuroR* and *NeoR* genes, the *D. melanogaster tra* female-specific intron (*traF*) located between *tra* exon 1 and 2 (3L: 16591003-16590756, 248 bases, FlyBase.org) or the *D. melanogaster dsx* male-specific intron (*dsxM*) located between *dsx* exon 3 and 5 (3R: 7930688-7935767, 5080 bases) was inserted inside coding sequences of antibiotic resistance genes. The plasmids generated in the study and their complete sequences (Figure 1a) were deposited at www.addgene.org (#131613 – #131618).

### Fly Transgenesis

The generated PB plasmids were injected into w^1118^ flies at Rainbow Transgenic Flies, Inc. (http://www.rainbowgene.com). Recovered transgenic lines were balanced on the 2^nd^ and 3^rd^ chromosomes using single-chromosome balancer lines (w^1118^; CyO/sna^Sco^ for II and w^1118^; TM3, Sb^1^/TM6B, Tb^1^ for III) or a double-chromosome balancer line (w^1118^; CyO/sp; Dr/TM6C, Sb^1^). Multiple independent lines integrated on the 2^nd^ and/or 3^rd^ chromosome were recovered for each plasmid and tested on food supplemented with puromycin or geneticin. We used heterozygous transgenic lines harboring one copy of a transgene to assess the antibiotic resistance and sex-sorting efficiency. One transgenic line harboring the complete sex-sorter cassette integrated on the 3rd chromosome supported 100% sex-sorting of both sexes, was homozygous fertile, and demonstrated especially good fitness. This line with two copies of the sex-sorter cassette (homozygous) was used for further analysis and deposited at Bloomington Drosophila Stock Center (BDSC #79015).

### Genetics and Sex Selection

Flies were maintained on the cornmeal, molasses and yeast medium (Old Bloomington Molasses Recipe) at 25□°C with a 12H/12H light and dark cycle. To assess drug resistance and/or sex selection, we used the Instant *Drosophila* Food (Formula 4-24) by Carolina biological Supply Company. A 1.1 g of the dry food was aliquoted per fly vial (FlyStaff.com) and mixed with 5 mL of distilled water supplemented with puromycin (Sigma #P8833) or geneticin (G418, Sigma #A1720), in varying concentrations from 0 to 1.2 mg/mL. To assess the drug resistance and/or gender sorting of transgenic flies, a mixture of *wt* and transgenic flies harboring one copy of a transgene were allowed to breed on the supplemented food (Figure 1c). Once the 3^rd^ instar larvae began to appear, the parents were removed, and after hatching of the adult offspring, their transgenic markers and gender were recorded. Two or three independent transgenic lines integrated on the 2^nd^ or 3^rd^ chromosome were analyzed on food supplemented with puromycin and/or geneticin at 0.4 and 1.0 mg/mL. Since puromycin is known to be unstable over time in water solutions, we counted emerging flies and noted their gender for only seven days after the first fly emerged. Three or five replicates were set up for each concentration.

### Fitness Estimation

To compare fly fitness on different food regimens, we calculated the percentage of embryos that survived to adulthood on the Instant Drosophila Food (Formula 4-24). Large numbers of *Drosophila* embryos were staged and collected on grape juice agar plates that were fitted into embryo collection cages (Genesee Scientific, FS59-100) following the started protocol. In short, 50–80 flies were transferred into embryo collection cages and laid many eggs on grape juice plates fitted on the bottom of a cage. Grape juice plates were replaced, and the embryos laid on the plates overnight were collected in the morning. Embryos of both *wt* and transgenic flies harboring two copies of the sex-sorter cassette were collected. Batches of seventy five embryos were placed on the instant food supplemented with 0.0, 0.4, and 1.2 mg/mL of puromycin or geneticin; and the number and gender of emerging adult flies were recorded for each condition. For the sex-sorter line raised on foods with different antibiotic concentrations, the embryo-to-adult survival rate of either males or females on drug-supplemented foods was compared to that of the corresponding sex for *wt* embryos developed on the food without any antibiotics. In other words, the survival percentage of each gender were compared and presented as normalized percentages (Figure 2d). The embryo-to-adult survival rate was estimated from five biological replicates. Finally, to assess the competitiveness of drug-sexed homozygous *Opie2-NeoR*^*traF*^+*Opie2-PuroR*^*dsxM*^ males (line #3, Figure S1d) against *wt* males, we placed one transgenic male raised on food supplemented with 0.4 mg/mL of puromycin and one age-matched *wt* male into a vial with ten virgin females and let me compete with each other for female mating. Their progeny was screened for the dsRed transgenic marker, and the percentage of their progeny sired by a homozygous transgenic male was calculated for ten independent vials. The mean (± SD) percentage of transgenic progeny was compared to 50%, which is expected when both males are equal at female mating.

### Drug Selection Stage

In order to determine the larval stage at which gender selection occurs, we fed first instar *wt* larvae with a yeast paste with or without drug at 0.4 mg/mL, and observed the larval stage to which they survived. *D. melanogaster* embryos were staged and collected on grape juice agar plates as for the fitness comparison. Then, the embryos were transferred on agar plates with a yeast paste supplemented with drugs spread on the surface, incubated at 25°C, and the endpoint of their development was observed and recorded.

### Statistical Analysis

Statistical analysis was performed in JMP 8.0.2 by SAS Institute Inc. The percentages of a specific gender or embryo-to-adult survival were compared with the corresponding values estimated for the *wt* flies (Figure 2a–c). *P* values were calculated for a two-sample Student’s *t*-test with equal variance.

## ASSOCIATED CONTENT

### Supporting information

**Table S1:** Development of sex-sorter system

**Table S2:** Titration of drugs for homozygous sex-sorter line

**Table S3:** Survival rate for homozygous sex-sorter line

**Table S4:** The competitiveness of males sexed by puromycin

## DATA AND REAGENT AVAILABILITY

All data that are represented fully within the tables and figures. The plasmids constructed in the study (Figure 1a) were deposited at Addgene.org (#131613 – #131618). The homozygous sex-sorter lines was deposited at Bloomington Drosophila Stock Center (#79015). The remaining *Drosophila* lines will be made available upon request.

## AUTHOR CONTRIBUTIONS

O.S.A conceived the idea, engineered plasmids, and carried out preliminary tests at Caltech. O.S.A and N.P.K designed experiments. N.P.K, A.D.H. and J.L. performed all molecular and genetic experiments. All authors analyzed the data, contributed to the writing of the manuscript, and approved the final manuscript.

## ACKNOWLEDGMENTS

This work was supported in part by NIH grants 5K22AI113060, 1R21AI123937 and UCSD startup funds awarded to O.S.A.

---

**Figure S1.**
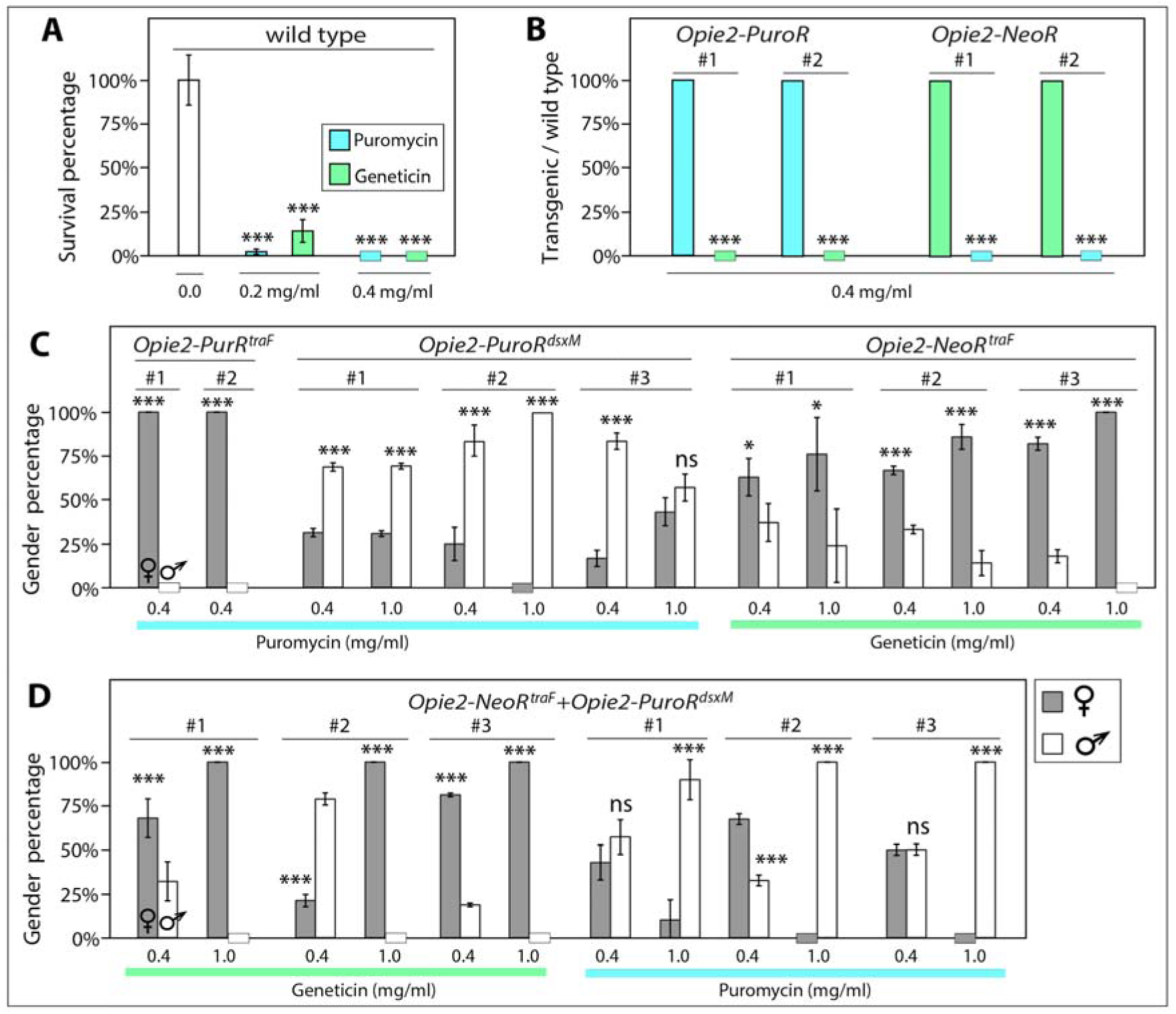
Development of sex-sorter cassette in *Drosophila*. (a) Supplementing fly food with puromycin or geneticin to the final concentration of 0.4 mg/mL completely arrests development of wild type (*wt) D. melanogaster*. Both drugs are also toxic to *wt* larvae at 0.2 mg/mL, but a few adult flies do emerge. (b) Fly survival in two independent transgenic lines harboring one copy of either *Opie2-PuroR* or *Opie2-NeoR* relative to *wt* flies on food supplemented with 0.4 mg/mL of puromycin or geneticin. The *PuroR* and *NeoR* genes expressed under the *Opie2* promoter rescue survival of transgenic flies harboring one copy of a transgene on the corresponding drug, while all *wt* flies perished. (c) The sex-specific drug resistance is achieved by inserting of the female-specific *traF* or male-specific *dsxM* intron into coding sequences of *PuroR* and *NeoR*. The efficiency of drug-induced sex sorting was assessed for a few independent transgenic lines of the same genetic construct. Transgenic flies harboring one copy of the antibiotic resistance genes were raised on drug-supplemented food. When sex sorting was not 100% efficient, the higher drug concetration of 1.0 mg/mL was used. (d) Both antibiotic resistance genes expressed in the opposite sexes were combined into one sex-sorter cassette, *Opie2-NeoR*^*traF*^*+Opie2-PuroR*^*dsxM*^. Three independent transgenic lines harboring one copy of the sex-sorter cassette were tested on food supplemented with either puromycin or geneticin at 0.4 and 1.0 mg/mL. We found two transgenic lines that can produce 100% males or 100% females when raised on food supplemented with 0.1 mg/mL of geneticin or puromycin, respectively. Bar plots show the average ± SD over at least three biological replicates. Statistical significance was estimated using a *t* test with equal variance. (*P* ≥ 0.05^ns^, *P* < 0.05*, *P* < 0.01**, and *P* < 0.001***).

**Figure S2.**
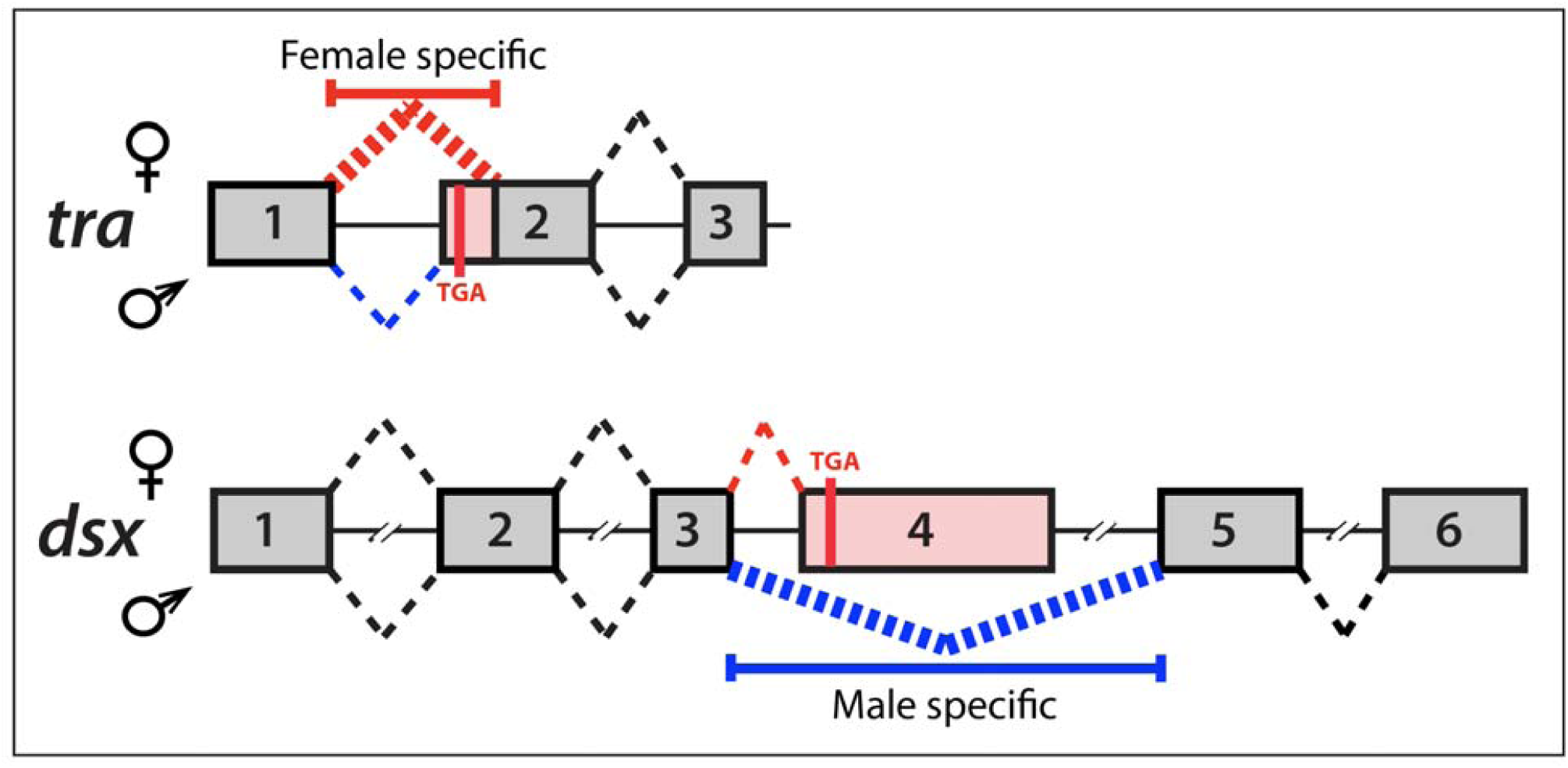
Sex-specific alternative splicing of *Drosophila double sex* (*dsx*) and *transformer* (*tra*) gene. The female-specific intron between *tra* exons 1 and 2 (*TraF*) is spliced out entirely in females, while in males some sequence remains producing a stop codon resulting in premature termination of Tra protein^43^. Inversely, the male-specific intron between *dsx* exons 3 and 5 (*dsxM*) is spliced out in males, while in females only its small part is spliced out and an entire exon 4 carrying a premature stop codon remains^44^. To establish female- or male-specific expression of antibiotic-resistance genes, we inserted *TraF* or *dsxM* intron sequences, correspondingly, into the coding sequences of *PuroR* and *NeoR*.

